# Reconstructing gene content in the last common ancestor of cellular life: is it possible, should it be done, and are we making any progress?

**DOI:** 10.1101/013326

**Authors:** Arcady Mushegian

## Abstract

I review recent literature on the reconstruction of gene repertoire of the Last Universal Common Ancestor of cellular life (LUCA). The form of the phylogenetic record of cellular life on Earth is important to know in order to reconstruct any ancestral state; therefore I also discuss the emerging understanding that this record does not take the form of a tree. I argue that despite this, “tree-thinking” remains an essential component in evolutionary thinking and that “pattern pluralism” in evolutionary biology can be only epistemological, but not ontological.

---

This essay discusses the problem of inference of gene content in the Last Universal Common Ancestor of all cellular life on earth (LUCA). The task of inferring an ancestral genic character is this. We have genome sequences of several currently existing species containing information about the states, such as presence or absence, of a homologous character – most often, a homologous fragment of the nucleotide or amino acid sequence – that is found in some or all of these genomes. We want to infer the state of that homologous character in the most recent common ancestor of these genomes. The algorithms that perform the inference are typically recursive, i.e., the states at all the intermediate steps between the present-day species and the common ancestor are identified in order to define (“retrodict”) the ancestral state.

Within this approach, different kinds of characters can be examined. For example, we may seek to retrodict the state of nucleotides, codons or amino acids within a set of homologous genes, to get at the identity of each character in the ancestral gene or protein sequence. Interestingly, the deduced sequence can be actually synthesized (“resurrected”), and its properties can be studied experimentally. In another kind of reconstruction, the trait of interest is the gene as a whole; we take as the input the list of orthologous genes found in the currently available genomes, to get at the status of the entire gene, such as its presence, absence, or perhaps copy number, in the ancestor.

Here I focus on the approaches that make inferences of the second type, i.e., examine genes from the extant genome sequences and retrodict the status of ancestors of these genes in LUCA. The task requires several inputs, most importantly: 1. a record of gene states in the extant completely sequenced genomes, together with the information about the orthology relationships between genes in different genomes; 2. a genealogy of genome lineages leading from LUCA to the present-day genomes, and of genes that evolve within these lineages; and 3. a description – best of all, a quantitative model – of the evolutionary process of gene gains and gene losses in the genomes.

The concepts of homology and of its special case, orthology, as well as the methods of computational definition of orthologs, have been reviewed extensively (Kristensen et al., 2011; Trachana et al., 2011; Altenhoff and Dessimoz, 2012; Sonnhammer et al., 2014). Several open questions notwithstanding, the field is approaching maturity, and various resources containing precomputed orthologous sets of genes are available, as are practical algorithms of *de novo* ortholog definition; all this will not be discussed here. Instead, I will examine the progress in the development of two other resources needed for reconstruction of LUCA gene content, i.e., the models of gene gain and loss in evolution, and the representation of the evolutionary history along which these gains and losses occur.

Thus far, I have evaded the question of what the genealogy of species actually looks like. For a long time, researchers presumed that such a history can be appropriately represented as a phylogenetic tree, i.e., as an acyclic graph. But in the last decade, with the accumulation of diverse, complete genome sequences and application of phylogenetic approaches at the genome scale, we learned that horizontal transfer of genes – the examples of which, of course, have been noted before completion of any full genome sequences – is a frequent, ongoing process in most lineages of Life, and therefore the history of Life has horizontal branches; it is not a tree in a precise sense. The apparently vast extent of horizontal gene transfer (HGT) in evolution poses a two-fold question in the context of LUCA reconstruction: If the history of Life is better modeled by something other than a tree, can we know what this “something” is? And can we retrodict anything on that non-tree-like history of Life?

To offer my opinion of these matters, in the rest of this paper I will have to switch directions twice – starting from the discussion of studies of the ancestral gene content problem when the genealogy is presented in a tree, without too much regard given to HGT events; moving on, to examine the re-evaluation of Tree of Life and recent proposals to replace it with something else, in the light of the role of HGT; and then back again, to the implications of all this for current and future research on LUCA gene retrodiction.

## Gene gain, gene loss, their relative rates, and retrodiction of the ancestral gene content

Gene gains and gene losses start with a molecular event in a cell. Addition of a new gene to a genome may occur in several ways, including: duplication of existing genes, followed by sequence divergence; gene/protein domain rearrangement; gene recoding, or overprinting, where a formerly non-coding, or coding in a different phase, segment of DNA becomes a part of a new gene; and, finally, HGT of a gene from either closely related or distant organism. Gene losses may occur by mutational inactivation followed by sequence deterioration, or by deletion of a large portion, perhaps the entirety, of a gene. Gene losses and gene gains can involve not only one gene at a time, but whole groups of genes. At the evolutionary scale, however, we are interested not so much in the rates of all those events in any particular cell, but in the rates of fixation of gene gains and gene losses in the populations of organisms, and ultimately in the whole lineages that we are examining. It is this fixation rate that is relevant for long-term gene retention and functioning of genetic systems in evolution, and here I speak of gene gain rate and gene loss rate in that sense.

Duplication of existing genes has been proposed, as early as in the 1960s, to be the major route of gaining new genes by the genomes, at least in vertebrates (Ohno et al., 1968), but the full extent of gene gain by duplication in any genome, and in the evolution of Life, could not be ascertained with partial genome sequences. Before genome-scale sequencing, we had even less confidence in gene absence, as one could always speculate that a homolog of any given gene resided in the yet-unseen portion of a particular genome. All this has changed in the 21^st^ century The genomic era provides us with a wealth of DNA sequence information from which past molecular evolutionary events can be inferred, often with a defined degree of confidence. One aspect of molecular evolution that has been illuminated by large-scale genome comparisons is the important role of ongoing gene gain and gene loss in the history of all genomes.

Given a collection of species with completely sequenced genomes, and a gene in one of these genomes, we can find the orthologs of that gene in all other genomes that have it; the set of such orthologs is sometimes referred to as “the same gene in different genomes”. Of course, some genomes may lack the ortholog of a given gene, so for each gene in any genome, its phyletic vector can be defined as a string of ones and zeroes that encodes the presence and absence of its orthologs – “the same genes” – in different genomes^2^. Sometimes a gene has several closely related, lineage-specific paralogs – in-paralogs – in the same genome. Such in-paralogs would be all orthologous (“co-orthologous”) to an ortholog in another lineage. Because of in-paralogy/co-orthology, the state of “one” (presences) in the phyletic vectors can be replaced by the actual counts of in-paralogs in each lineage. Thus, gene phyletic vectors capture information about sets of orthologs and co/orthologs, as reflected in a seemingly-redundant name of the first orthology resource, NCBI COGs – Clusters of Orthologous Groups (Tatusov et al., 1997). In the following, I use “COGs” in the generic sense of any collection of orthologous groups, unless a specific study is cited that utilized a particular version of NCBI COG resource.

Since most genes have orthologs in only a subset of completely sequenced genomes, phyletic vectors of most genes contain at least some zeroes. In fact, only a small proportion of all genes/COGs – less than 50 genes by the current account – are found in every sequenced genome without exception. From the evolutionary point of view, an explanation for a vector with all coordinates set to one is that the genes that have such a vector were present in LUCA and were strictly vertically inherited by all its ancestors, including the extant species with completely sequenced genomes. This is explanation is often, though not always, correct.

There are also COGs that are found in *almost* all species: for example, several dozen COGs are found in 95-99% of all completely sequenced genomes (Mushegian, 2008). An intuitive, and also often correct, explanation in this case is that such COGs were found in LUCA and have been preserved in the majority of lineages by vertical descent, and lost only in a few of them.

Matters become more complicated when we examine the remaining majority of COGs. Those are distributed sparsely. Consider the simple inference just described, i.e., “a gene was present at the root of the tree and was occasionally lost on the way to some of the present-day species”. This looks like a straightforward scenario for the COGs that are found in the majority of species, but it faces a difficulty when applied to the whole set of phyletic vectors. Examination of one version of prokaryotic COG resource has shown that 90% of all COGs are found in 20 or fewer species out of total 110 (Mushegian, 2008). An attempt to explain each such sparse vector only by losses would be the same as assuming that gene losses are more common in evolution than gene gains by almost two orders of magnitude; each of these COGs would have experienced one gene gain at the root of the Tree of Life and more than 90 gene losses in their entire history. Such a scenario would mean also that no new genes have emerged since LUCA, and that the LUCA genome contained the ancestor of every COG, resulting in a large genome size of 14000 genes. None of this is supported either by common sense or by more specific comparative-genomic considerations (see below), and no model that I know of uses this estimate for LUCA retrodiction^3^. This did not prevent W. F. Doolittle from preposterously purporting in his essay (Doolittle, 2009) that my earlier discussion of LUCA (Mushegian, 2008) suffers from the wrong assumption that LUCA contained the ancestor of every present-day gene (whatever that means).

The model of “the mother of all genomes evolving by too many gene losses” may be partially remediated by the fact that many of the COGs are found only within a monophyletic subtree of the species’ tree. Phyletic distribution of such COGs may be explained by gene gain in the last common ancestor of the species that currently have this COG, i.e., in a species that is represented by an internal node in the tree, not by LUCA. This reduces the excess of gene losses over gene gains in each phyletic vector, because a gene is gained once at the root of the subtree, and no losses are invoked to explain its absence outside the subtree. This also provides a more plausible picture of gene emergence – different genes first appear in the genomes at different times. Such a scenario is incorporated in any evolutionary model of gene content but, as discussed below, the models that include only gene losses turn out to be inferior to the models that also allow gene gains in the course of evolution of each orthologous group.

The seminal work by Mirkin and co-authors (2003) presented the algorithms to infer the ancestral state of each COG by determining the “amount of evolution” explaining each phyletic vector, given the species tree. Proceeding from the currently known states of genes at the tips of the phylogenetic tree, it attempted to reconstruct the state of each COG at each intermediary node of the tree. The approach utilized a weighted parsimony criterion – for each phyletic vector, such set of changes was selected that minimized the score *S* = *λ* + *gγ*, where *λ* is the number of losses that occurred during the evolutionary history of a COG, *γ* is the number of gains during that history, and *g* is the “gain penalty parameter”.

Assume that each gene observed in the data at the very least had experienced one gain event, upon its first appearance. Therefore, *γ* is always 1 or more, *λ* is zero or more, and the value of *g* can vary. Intuitively, for positive *γ* and *λ*, when *g* >>1, the score will be dominated by the second term – many gene losses will count the same as one gene gain. In this case a phyletic vector with plenty of zeros must be explained mostly by gene losses, which are relatively cheap. This places the first emergence of a gene back in time, closer to LUCA. Conversely, when *g* <<1, many gene gains count the same as one gene loss, and to minimize the score, vectors tend to be explained by including more gene gains.

Mirkin et al. explored different values of gene gain penalty and asked, for each value of *g*, what the gene complement in LUCA would look like. Most interestingly, at *g* ≈1, the sum of all *S* values explaining each observed phyletic vector in the COG database was at a deep minimum. The distribution of COGs by the number of events explaining their evolution in that case had a peak around 3, suggesting that most genes may have experienced ≤2 events in addition to their birth. In other words, a substantial fraction of genes may have been horizontally transferred either once or never in their history, and a minority of genes – though in total, this is still a large number – must have been transferred more frequently. The average of three events, one of which is a gain, gives at most two-fold excess of losses over gains.

Examination of the molecular functions of genes retrodicted into LUCA when the value of *g* was set at 1 (called LUCA1.0, with 572 COGs) indicates also that this assembly of genes, constructed without taking any functional information into account, is nonetheless biochemically coherent. LUCA1.0 encoded not only the nearly-full complement of the components of RNA translation apparatus (this cellular module is consistently inferred by all models of LUCA inference, though important details vary between the models), but also substantial portion of the core intermediary metabolism. In particular, glycolysis and de novo biosynthesis of nucleotides were retrodicted in nearly complete form, and parts of amino acid biosynthesis was also present.

The evolutionary model of Mirkin et al. was based on weighted parsimony, counting and weighting the events in steps between each node in the tree, accounting implicitly for different probabilities of gene gains and gene losses. Direct ways to introduce probabilities are better, and more recent inferences of LUCA gene sets have been done with probabilistic, maximum-likelihood models. In one study (Cohen et al., 2008), the joint averages for “gene gains/losses” (GGL) rates were reported, reaching about 7 in the most parameter-rich models; this value, however, is influenced by the subset of large families with extremely high GGL, whereas more than half of all phyletic vectors have GGL ≤ 3.

Estimates of the gene loss and gain ratios also can be done on the datasets that represent individual clades, to learn about immediate ancestors of that clade. Some of these studies, though not dealing with LUCA directly, provide us with useful insights about the parameters of the process. For example, a maximum-likelihood approach in the context of a gene birth-and-death model has been applied to the study of gene content in 28 genomes of archaea (Csurös and Miklós, 2009). Gain by horizontal gene transfer and gain by gene duplication were modeled as two separate processes, variation of these rates between families was accounted for, and likelihood computation was improved by correcting for the complete extinction of some genes. The average loss-to-gain ratio across all families and all lineages was close to 2. However, gain rates in the individual lineages were spread twice as wide as loss rates, and in some lineages the rates of gains, losses and duplications were all far from average (indeed, a handful of lineages were dominated by gene gains). The ancestral genome size, at ~1200 genes, was slightly smaller than the genomes of most extant archaea.

More extreme views of the loss-to-gain ratios and LUCA genome sizes are also known. For example, a study in a high-profile journal (David and Alm, 2011) described a birth-and-death model of gene content evolution that gave a tiny gene count in LUCA (below 200). Though many technical details of the methodology employed in that study remain unclear, it appears that the model works out to about 0.3 gene losses per one gene gain averaged across all lineages. On the other end of the opinion spectrum is an assertion of the loss-to-gain ratio in bacteria being about ten (Cavalier-Smith, 2002), which does not cite any compelling evidence, but, as far as I can discern, would set the size of the bacterial ancestor at ~9,000 genes.

All the efforts that I described thus far rely on the existence of the universal history of Life, and moreover depend on the Tree of Life as the true record of that history. Horizontal gene transfer is acknowledged in these approaches, but is viewed mostly as the difference between in the fate of an individual gene and the topology of the independently objective Tree of Life. But the truth of this representation has been brought into doubt recently by the results of comparative analysis of completely sequenced genomes, and in particular by the understanding of widespread horizontal gene transfer in the evolution of Bacteria and Archaea. The corollary of those recent findings is the idea that the history of Life on Earth should not be represented as a Tree. And to some in the research community, a corollary of such a corollary seems to be that reconstruction of LUCA is a futile exercise – definitely in our current state of confusion about the history of life, and perhaps even in principle. In the next section, however, I would like to argue that, while the corollary is true, the corollary of the corollary is false: the history of life is Not A Tree, but it may be recoverable just the same, albeit as another kind of a graph. And inference of the ancestral states may be possible on such a graph, too.

## A Tree or not a Tree, and so what?

The evolutionary history of cellular Life on Earth cannot be represented as a tree in the computer-science sense, because it contains cycles (reticulations or horizontal branches), which represent various evolutionary events such as horizontal gene transfer and possibly also whole-genome mergers. This became evident decades ago, when the hypothesis of endosymbiotic origin of mitochondria and plastids (Wallin, 1925, Sagan, 1967) started to receive confirmation from comparative sequence analysis (Steinman and Hill [1973] may have been the first to provide the sequence-based evidence). The new idea for the 21^st^ century is that molecular and computational methods can be applied to detect and quantify many HGT events throughout the history of Life. Much work is being done to address this important conceptual and practical problem, and many basic parameters of HGT are now beginning to be understood. The emerging picture is complex: HGT has both “frequent” and “rare” aspects. Before going into this, let us review the primary evidence, approximately in the order of its appearance in the literature and in the consciousness of biologists.

First, an early “smoking gun” of HGT was the observation of highly similar genes encoded by various plasmids and integrated mobile elements in bacteria, some of which may be selfish genetic elements, but others of which provide the host cells with evolutionary advantages, such as detoxification of antibiotics, resistance to phages, or novel metabolic functions. Second, this is closely paralleled at the nucleotide level, as many such regions (as well as other, more “normal-looking” genes) have nucleotide frequency properties different from the averages for the the host cell. Third, as said already, there is plenty of cytological and molecular evidence that eukaryotes have acquired two large batches of bacterial genes by symbiogenesis: ancient alpha-proteobacteria have become mitochondria after being engulfed by an ancestor of eukaryotes, and additionally, probably more than once, different eukaryotes have acquired ancient cyanobacteria, which became plastids. Fourth, many pairs of distantly related microorganisms living in the same habitat tend to share more orthologs than non-cohabiting species from the same pair of lineages. Fifth, given a set of species and a set of genes shared by these species, trees of different genes from these species often have different topologies. In many of these cases, the known tree inference artifacts can be ruled out, and the incongruence must be explained by the different history of the genes in each tree, including differential HGT.

These kinds of evidence are not all the same. The first two groups of observations get the closest to the understanding of actual molecular mechanisms of gene transfer. They are special also because the inference of phylogenetic trees did not play a major role in the discovery of these phenomena, and they can be analyzed with little recourse to the trees. In contrast, the other lines of observation rely on the phylogenetic inference. Thus, the vast comparative-genomic evidence of HGT is invoked in the first place through examination of phylogenetic trees, albeit the trees of individual genes^4^. This is worth remembering while assessing the uses of trees and “tree-thinking” in biology (see below).

With the preponderance of this data, the questions about HGT are no longer existential, but rather mechanistic and quantitative ones, i.e., those concerned with the parameters of the HGT processes in nature. HGT can be examined in the context of gene families or in the context of genomes, at different taxonomic and temporal scales – and depending on the question of interest, the answer may come out on the “rare” or “frequent” side. For example, the average number of HGT events over the lifetime of an individual gene may be low – for most genes, <2 transfers over their entire history have been detected (Lerat *et al*., 2005; Creevey et al., 2011). On the other hand, the number of genes/COGs that have been horizontally transferred during their lifetime is high: a large fraction of COGs, amounting to many thousands of them, has been transferred at least once during their evolution (Koonin et al., 2001; Kloesges et al., 2011). Thus, “HGT is rare” and “all HGT, all the time” camps may in fact be representing the facets of a complex phenomenon, and it is likely that different views of HGT may be reconciled if a proper account is taken of what is being measured in each case.

Similar kind of careful parsing should be applied to the notion of “The Tree of One Percent”, which in its most direct form states that the “consensus” phylogeny of prokaryotes, built in the last decades of this century on the basis of universal molecular sequences such as ribosomal RNAs, is in fact not supported by phylogenies of other genes: if trees for all protein-coding genes are also examined, only a small fraction of them have the same topology as the rRNA-based trees (Dagan and Martin, 2006). This, the story goes, makes the rRNA-based phylogeny un-representative of the evolution of Life – it focuses on just one cellular subsystem, and ignores the majority of other evidence.

Another view, however, is that the tree first derived on the basis of rRNA is in fact a fully relevant representation of an essential evolutionary reality. The argument here is two-fold. First, analysis of ribosome composition, mechanism of its maturation, and molecular function shows that the genetic complement required for these omnipresent roles is not limited to genes encoding ribosomal proteins and RNAs; it also includes the enzymes involved in post-trancriptional modifications of rRNA and tRNA; in post-translational modifications of ribosomal proteins; in co-translational maturation of the protein products of translation; and, at a slightly longer metabolic distance, the systems for biosynthesis of biochemically diverse cofactors required for the aforementioned modifications, as well as the systems providing the amino acid substrates for protein biosynthesis. Thus, protein synthesis machinery is not a small and segregated handful of essential genes, but a module integrated into a much larger network of other molecular functions.

The second consideration deals directly with the topologies of trees that are different from the “ribo-tree” and are seen as the rebuttal of “ribo-centric” phylogeny. Here, as with the “A Tree or Not a Tree?” debate, the main question is misleadingly presented as a binary choice (“Same or Different Tree?”). In reality, however, the gene trees that are topologically different from the “Tree of One Percent” are not randomly distributed in the space of all possible trees; typically, two gene trees of a prokaryotic data set are much closer to each other than two random trees (this has been pointed out, among others, by Nicholas Galtier in his open peer review of Bapteste et al. (2009)). Thus, “being not the same” as the ribo-tree is neither here nor there; the real question is whether or not the ribo-tree is a major central trend in the space of trees, and whether the majority of all gene trees with their different topologies are located within a short distance from that tree. The answer to that question seems to be in the affirmative (Koonin et al., 2011; Puigbo et al., 2012).

Tree rejectionists have a different take on the problem. Their research program advocates “pattern pluralism”, which states that phylogenetic tree is just one of many possible structural patterns representing the evolution of life on Earth. From a resent manifesto of that program (Bapteste et al., 2009): “With regard to the tree of life, the pluralistic position has thus been regularly advanced by microbial phylogeneticists who have emphasized the diversity of evolutionary processes and entities at play in the microbial world […]. This group prefers to model evolution as a diverse set of processes acting on the histories of diverse kinds of entities generating, finally, a diversity of overlapping and cross-cutting patterns, corresponding to different evolutionary outcomes. For such pluralists, depending on the approach taken (e.g., the choice of sequence, the choice of the reconstruction method, the taxa of interest), a different evolutionary pattern may be generated (e.g. a reticulated network rather than a vertical tree)”.

Let us examine this quote further, and ask what is pluralistic about the approach summarized in it, and how these pluralities contrast with the more traditional research programs. It seems that three main things are plural: first, it is “the diversity of evolutionary processes and entities at play in the microbial world” – presumably, as opposed to more monistic understanding of such processes and entities by the evolutionary biologists in the last century; second, it is “a diversity of overlapping and cross-cutting [evolutionary] patterns”, presumably, again, as opposed to the unity, or at least, lesser diversity of patterns known by the evolutionists of yesteryear; and, third, corresponding to these diverse patterns, there are “different evolutionary outcomes”. The first two kinds of pluralities seem to be well within the realm of any developing science – our understanding of natural processes may become more nuanced, and be represented in a more complex form than before; we are just at such a stage in evolutionary biology now. It is the third aspect, of the “plurality of outcomes”, that seems to be the most radical departure from the earlier phylogenetic narratives.

Among the philosophical foundations of the pluralistic approach to phylogeny, its proponents (Bateste et al., 2009; Doolittle, 2009) prominently cite the eliminative pluralism, which has been developed by M. Ereshefsky in his work on the concept of prokaryotic species. The essence of Ereshefsky's proposal was to eliminate the term "species" in microbiology and to replace it with a plurality of more appropriate terms (Ereshefsky, 1992). He is explicit about the fact that his argument for pluralism is ontological: species, he argues, do really exist as different kinds of entities in nature, and not merely so seem to us because of our temporary lack of information about the world. This is where the “pattern pluralists” of phylogenetics in fact depart from Ereshefsky's eliminative pluralism. The fact of the matter is that there can be no ontological pluralism in reconstructing the history of Life, for an obvious reason: only one such history existed on Earth, therefore only one account of that history must be correct. Epistemological pluralism, however, is possible in this case also: different kinds of graphs, or perhaps also other kinds of representations, may model various aspects of evolution, or represent various snapshots of our still-incomplete knowledge of that unique history.

In this debate on philosophical pluralism, it may be worth remembering that the goal of the study of evolution is neither “to build a tree” nor to “produce the plurality of representations”. Those are just the means to the real and important goal: to understand what essentially happened^5^ during the span of evolution of Life.

As we are looking for a more detailed and realistic reconstruction of the history of Life, which would, in all likelihood, require complex graphs which Are Not Trees, it can be stated with some confidence that the habit of “tree-thinking” in evolutionary biology, as well as the use of computer-generated trees, will remain essential ingredient of our work in practical evolutionary genomics and phylogenetics. Most obviously, trees are the most appropriate form in which to represent the one process that undoubtedly was important in all domains, kingdoms or empires of Life, without exception – the process of descent, by duplication with modification, of DNA and its encoded products from ancestral DNA (the same is true for phylogeny of RNA genomes of viruses, of course). Further, “ribo-tree” is a faithful representation of the history of ubiquitous gene ensembles, central for all cellular life.

Even if the history of a group of genomes is not exactly a tree, nevertheless its model in the form of a tree can serve as a sensible, robust null hypothesis in any evolutionary analysis, subject to severe tests (Mayo, 1996), i.e., tests such that if the hypothesis is wrong, our probability of detecting this is likely to be high. This point is worth emphasizing, as it is precisely by comparison to “the tree hypothesis” that any evidence of horizontal gene transfer is usually detected in practice. Finally, new algorithms that are devised to detect complex relationships between entities, where hierarchies (i.e., tree-like structures) and reticulations coexist, usually include some steps where trees are constructed or evaluated.

These reasons to hold on to trees and tree-thinking are what J. Velasco calls “the modeling defense of trees”, i.e., notion that the trees are useful to scientists because they are good models (Velasco, 2012). He distinguishes several kinds of models (“idealizations”), such as minimalist, or Aristotelian, idealizations, when only a small subset of relevant causal factors is examined; Galilean idealizations, when deliberate distortions of reality are introduced for technical tractability reasons, and then removed at a later step; and, finally, multiple-models idealizations, when a series of possibly incompatible models, each with its own trade-off in representing reality, are constructed without a requirement of later reconciliation between them. All this seems to be a different formulation of epistemological pluralism discussed above. Even so, inference and analysis of phylogenetic trees remain important parts of an evolutionary biologist's accoutrement.

A much stronger defense of tree-thinking in evolutionary biology, however, is the fact that the ribo-tree “of one percent” is not merely a useful fiction, but in fact a clear and important central trend in the evolution of life; it has the topology to which a large proportion of other genes, notably those that are present in the majority of lineages, are statistically close (Puigbo et al., 2012). No one in their sound mind would dismiss the mean of a statistical distribution for its failure to be equal to all values of the variable simultaneously. Likewise, ribo-tree is just one tree in the distribution of different gene trees, but it plays a central role in our attempts to understand this distribution as the way of interrogating all gene and genome trees, in order to make them reveal the evolutionary signal.

The future algorithms for phylogenetic inference will be devised to reconstruct both vertical and horizontal transmission of genetic elements, and – returning to our main subject – the algorithms for inference of the ancestral states of the presently known genic characters will operate on the graphs that are more complex that the cycle-free binary trees. Thus, the new view of the History of Life on Earth will contain, within itself, the old view as a special case. One can consider this either a rejection of The Tree of Life, or else its generalization. The latter “generalizationist” or “accumulationist” account of the History of Life would not be dismissive of the discoveries of the earlier generations of scientists, but would integrate them with more precise findings of our times. Such an approach would be graceful, morally serious and, I suspect, give factually correct results. With this aspiration, we return to the summary of the recent work on LUCA.

## What is being done on gene retrodiction, and what remains to be done?

A probabilistic retrodiction of gene content in LUCA, which employed the maximum-likelihood approach, was published recently by Kannan et al. (2013). One novel ingredient of that work was to compute the probability, for each COG, that its ancestor was present in LUCA (“probability of a gene being ancestral”, or “*gene ancestrality*”). Another extra step was to estimate the rates of gene gains and gene losses from the data, using phyletic vectors that allowed more than one in-paralog in a species (so that gene gain 0→1 and gene duplication 1→*m* events could be modeled separately, and analogously for gene losses). Such more complex models turned out to assign relatively high ancestrality to rare genes, and to place them into the common ancestor, more often than simple models did. Notwithstanding this difference, the gain penalty parameter was estimated in all models to be between 2 and 4. A series of LUCA gene sets was generated, corresponding to different inference modules and at various cutoffs of “gene ancestrality”; one retrodiction, called LUCA ML 0.7 (i.e., ancestrality of each gene ≥0.7), had 571 genes and was the closest in size to LUCA 1.0 from Mirkin et al., (2003).

This most detailed ML-based reconstruction of LUCA thus far leaves a lot of room for improvement. For example, heterogeneity of the rates of gene gain and gene loss in different branches of the phylogeny was not modeled, and genes completely lost from all lineages were also not accounted for. Doing so in the future would improve the estimate of the number of genes in LUCA, though of course the identity of the lost genes would not be known and must be deduced in other ways.

Despite different assumptions, methods and datasets, there seems to be an agreement on several aspects of retrodiction. In particular, if we omit the most extravagant estimates, the majority of studies put the ratio of gene loss and gene gain rates close to 2, and the LUCA gene count is estimated to be between 500 and 1000-2000. Of course, the several-fold difference in the number of genes has profound implications for the metabolic capacity and other biological properties of LUCA; moreover, different estimates may give similar counts of genes in LUCA but differ in the identity of retrodicted genes, with the same effect.

Complementing these investigations is significant progress made recently in the area of algorithmic and statistical approaches to inferring HGT from molecular data. The main innovations of the last decade include: better understanding of measures of dissimilarity between trees (Bordewich and Semple, 2007; Boc et al., 2010; Kannan et al., 2011); algorithms for “reconciling” trees of different topology, for building phylogenetic networks, and for identifying donors and acceptors of horizontal gene transfer (Lin et al., 2006, 2007; Abby et al., 2010; Cohen and Pupko, 2010; Boc et al., 2012; Stolzer et al., 2012; Patterson et al., 2012; Layeghifard et al., 2013; Steel et al., 2013); application of bootstrap to assign confidence to the horizontal branches in phylogenetic networks (Boc et al., 2010); and algorithmic account of the incomplete taxon sampling in the data vs. the need for donor and acceptor to be each other's contemporaries (Linz et al., 2010; Szöllosi et al., 2013). There must be little doubt that these and other developments, together with the increasingly dense sampling of genomes and gene families by new-generation sequencing approaches, will provide us, in not-so-distant future, with a detailed picture of reticulate phylogeny of Life, and with the practical ways of inferring gene ancestral states on the networks of complex topology.

The question of the root position in the historic account of life on Earth is of special importance. Similar to the case of “tree-thinking”, the two-pronged argument can be made also for “root reasoning”. On the one hand, constraining the position of ancestral nodes, such as LUCA, improves our ability to make inferences about the past. If the parameters of evolution are known, we may explore the consequences of varying the root position and optimize it under chosen criteria. This is the “modeling defense of the root”. On the other hand, it is important to remember that not only the “ribo-centric” trees, but also the genetic code of all life are essentially the same. The current, nearly-universal code may have been preceded by multiple, possibly co-existing, genetic codes, but a population of genetic elements is likely to have attained a measure of functional complexity prior to “genetic annealing” and the emergence of essentially modern-type code (Woese, 1998; Vetsigian et al., 2006). Thus the present-day life is ontologically monophyletic, and the concept of “Rooted Net of Life”, reflecting this, has been advocated in the literature (Williams et al., 2011). Interestingly, given the phyletic vectors of the existing genes and a model of gene gain and loss, the counts of genes in LUCA are not very sensitive to the placement of the root within the Tree of Life (Dagan and Martin, 2007). The identity of the ancestral genes, of course, would depend on placing the root correctly.

Lastly, it is important to note that the work of LUCA reconstruction is not done when the ancestral gene set is retrodicted. On the contrary, at that stage the reconstruction can be subject to testing, using several independent criteria, on which the inference did not depend. First, as already mentioned, the ancestral gene repertoire can be evaluated for the internal coherence, i.e., whether the set of metabolic pathways, precursor, products and gene fluxes makes functional sense (Mirkin et al., 2003; Gil et al., 2004; Tsoy et al., 2013). Second, the LUCA genome and the properties of its putative encoded proteins have to be evaluated in view of geology and biogeochemistry of the likely ancestral habitats – an area of study that itself undergoing profound transformation and rapid development in this century (Martin et al., 2008; Mulkidjanian et al., 2012). Third, much of this information will be tied together as we are improving the methods of resurrecting ancestral sequences, and studying the properties of single proteins and eventually protein ensembles from LUCA in the test tube (see Groussin et al., 2014 for recent discussion of technical issues and Gaucher et al., 2008, Zhou et al., 2012; Akanuma et al., 2013; Risso et al., 2014, for examples of resurrection at the LUCA-relevant time horizon).

The successes and failures of LUCA reconstruction will be likely recorded not only in individual research papers, but also in LUCApedia (Goldman et al., 2013).

## Footnotes

2 Gene phyletic vectors are also known as “phylogenetic patterns” or “phylogenetic profiles”. I prefer “vector” because this is what the construct is, mathematically speaking; in fact, computations on these vectors using linear algebra techniques are appropriate for answering various questions of biological interest, though this is a story for another day. “Phyletic” is also better than “phylogenetic”, because information about phylae, i.e., the tree tips, is explicit in the vector, whereas information on phylogeny is only implicit.

3 The other extreme in modeling the evolution of gene content is to postulate that no gene is ever lost from any genome. As far as I know, this class of models has not been explored systematically, probably because the falsity of such an assumption is quite self-evident.

4 One may object, of course, that many of the same inferences can be made by the analysis of nearest sequence neighbors in the context of sequence similarity search, without ever constructing a tree. This indeed is the way in which such analysis is often done in practice, but the legitimacy of that surrogate approach comes from the fact that under certain conditions, high-ranked pairwise similarity is a good predictor of the phylogonetic relationship between two sequences; despite some skepticism (e.g., Koski and Golding, 2001), and more recently validated quite extensively, e.g., in the context of ortholog definition (Altenhoff and Dessimoz, 2009).

5 The dictum by Leopold von Ranke (*wie es eigentlich gewesen*) had been translated into English at some point as “how things actually were” and criticized from various points of view, mostly in connection with scientists' inability to reconstruct the past fully and precisely, free of their own biases. However, several scholars, for example Bebbington (1979) have argued that a more accurate translation is “how things essentially were”, and this seems to be a good standard to which one may hold any historic reconstruction.

